# Multiple resource use strategies confer resilience to the socio-ecosystem in a protected area in the Yucatan Peninsula, Mexico

**DOI:** 10.1101/2020.01.08.897462

**Authors:** Luis Guillermo García Jácome, Eduardo García-Frapolli, Martha Bonilla Moheno, Coral Rangel Rivera, Mariana Benítez, Gabriel Ramos-Fernández

## Abstract

Natural Protected Areas (NPAs) are the main biodiversity conservation strategy in Mexico. Generally, NPAs are established on the territories of indigenous and rural groups driving important changes in their local resource management practices. In this paper we study the case of *Otoch Ma’ax Yetel Kooh*, an NPA in the Yucatan Peninsula, Mexico, that has been studied in a multidisciplinary way for more than twenty years. This reserve and its buffer zone is homeland to Yucatec Mayan communities that until recently used to manage their resources following a multiple use strategy (MUS), which involves local agricultural practices and has been proposed as resilience-enhancing mechanism. However, due to the restrictions imposed by the decree of the reserve and the growth of tourism in the region, some of these communities have started to abandon the MUS and specialize on tourism-related activities. We build a dynamical computational model to explore the effects of some of these changes on the capacity of this NPA to conserve the biodiversity and on the resilience of households to some frequent disturbances in the region. The model, through the incorporation of agent-based and boolean network modelling, explores the interaction between the forest, the monkey population and some productive activities done by the households (milpa agriculture, ecotourism, agriculture, charcoal production). We calibrated the model, explored its sensibility, compared it with empirical data and simulated different management scenarios. Our results suggest that those management strategies that do not exclude traditional activities may be compatible with conservation objectives, supporting previous studies. Also, our results support the hypothesis that the MUS, throughout a balanced integration of traditional and alternative activities, is a mechanism to enhance household resilience in terms of income and food availability, as it reduces variability and increases the resistance to some disturbances. Our study, in addition to highlighting the importance of local management practices for resilience, also illustrates how computational modeling and systems perspective are effective means of integrating and synthesizing information from different sources.

## 1 INTRODUCTION

Natural Protected Areas (NPAs) are one of the main conservation strategies in the world to face biodiversity loss. In 2016, 14.7% of terrestrial and inland water ecosystems worldwide were under this type of protection (UNEP-WCMC and IUCN, 2016). This strategy, however, has been highly controversial for their impact on the rural communities that inhabit the protected territories, namely, on their forms of resource management and appropriation (Durand and Jímenez, 2010; García-Frapolli, 2015). Nowadays, conservation strategies in NPAs range from total exclusion of human activities, to the promotion of the sustainable use of the territory (Brockington and Wilkie, 2015).

Mexico, as one of the biologically megadiverse territories in the world, has adopted NPA as its main governmental conservation strategy. Nationwide, 10.87% of the terrestrial and inland water ecosystems are under protection (CONANP, 2018). In addition, the country also harbours a great cultural diversity and indigenous groups territories generally are on zones of high biodiversity (Toledo, 2013). Therefore, many of NPAs in Mexico are inhabited by indigenous communities with long-lived productive and cultural practices, and as such, they constitute a highly diverse and complex social-ecological systems (SESs). SESs are indeed conceptualized as complex systems in which humans are considered as part of nature (Berkes et al., 2003). In contrast to traditional ecological and social research, studies with a SES perspective explicitly consider ecological and human components and their interactions (Liu et al., 2007). Additionally, in SES research there is a particular interest in studying and understanding feedbacks, nonlinear dynamics, thresholds, unintuitive behaviors, time lags, heterogeneity and resilience (Liu et al., 2007).

As the world undergoes unprecedented global changes, SES research has been particularly interested in understanding resilience and characterizing resilience-enhancing mechanisms (Biggs et al., 2012). This quest has lead to recognizing the value that some traditional or local practices, largely overlooked and blamed as unproductive and environmentally damaging, may actually have for resilience (Berkes et al., 2000). An example of these is the multiple-use strategy (MUS) on which indigenous communities often base their resource management and family household (Toledo et al., 2003). MUS involves the development of a set of different productive activities (e.g., agriculture, agroforesty, gathering, etc.) on a diversity of land units (e.g., milpa, successional forest, mature forest, etc.). It has been argued that this strategy amplifies the subsistence options available for the households to ensure a continuous flow of goods and services, thus minimizing the vulnerability associated with different disturbances (Barrera-Bassols and Toledo, 2005). However, integral assessments of coupled environmental and social outcomes of MUS are largely lacking.

In this paper we study the MUS practiced by a Mayan community inhabiting *Otoch Ma’ax Yetel Kooh* (OMYK, “house of spider monkey and jaguar” in Yucatec Maya), an NPA located at the northeastern part of the Yucatan Peninsula, Mexico. This NPA was declared in 2002 in response to a local initiative, after years of community-based conservation. It consists of a body of lakes surrounded by forest in multiple successional stages that harbours a large population of spider monkeys and other ecologically important species. OMYK and its buffer zone are inhabited by Yucatec Maya communities, which, until recently, used to manage their resources following a MUS based on traditional swidden milpa agriculture (García-Frapolli et al., 2008). The milpa system consists of a polyculture which typically combines maize, beans and squash, along with other domesticated, semi-domesticated and tolerated species (Benítez et al., 2014). Milpa system is highly diversified and adapted to an ample variety of environments, playing a key role in the maintenance of biological diversity and food sovereignty (Altieri et al., 2012). However, due to the growing tourism industry in the region and the restrictions imposed by the decree of the NPA, the local communities have experienced important changes in their management strategies. In fact, some households from the community within the NPA have shown a trend towards abandoning their MUS and specializing in the provision of ecotourism services (García-Frapolli et al., 2012; Rios-Beltrán, 2016).

These important changes lead us to ask the following questions: is the elimination of the traditional management practices, particularly the milpa agriculture, the only way to ensure the conservation of biodiversity in OMYK? And, how do different productive strategies affect the resilience of this SES in face of some typical disturbances of the region (e.g. hurricanes, fires, tourism-related fluctuations)? These questions have been previously explored in separate works, focusing on either environmental or social aspects. García-Frapolli et al. (2007), using a probabilistic model, proposed that traditional activities are compatible with conservation objectives in the reserve. In another work, García-Frapolli et al. (2012) have proposed that the MUS is a mechanism that promotes household resilience as it diversify the subsistence means. In this work, we address these questions together, using an integrative and dynamical computational model, a novel approach to understand the SES.

Dynamical computational models allow us to make reproducible experiments in fully controlled virtual SESs, on relatively short periods of time and without directly affecting the people that inhabit them (Barreteau et al., 2001). Agent Based Models (ABMs) are a type of dynamical model that has been widely used in the study of SESs (An, 2012). In ABMs a system is modeled as a set of individual interacting elements, or agents. Each agent owns a set of variables that describe its state and a set of rules that determine its behaviour (Railsback and Grimm, 2012). Some examples of the use of ABM to study SESs include the study of ecological degradation scenarios on indigenous communities on Amazonian Guyana (Iwamura et al., 2014, 2016); the study of the impact of demographic human changes on the deforestation of the panda habitat on the Wolong reserve, China (An et al., 2005); and the study of the effect of different management strategies on “La Sepultura” reserve, Mexico (Braasch et al., 2018). Boolean Network Models (BNMs) are another type of dynamical models that provides a qualitative representation of a system. BNM are discrete models in which a system is represented as a directed graph whose nodes represent the variables of the system and edges represent regulatory relationships between them (for a detailed description see Saadatpour and Albert (2013)). BNM have been mainly used in the study of molecular and cellular scale systems and it has not been until recently they have been used to study ecological and agroecological systems (e.g. Robeva and Murrugarra (2016); Gaucherel et al. (2017); López-Martínez (2017)).

The objective of this study is to explore the effect of different productive and management strategies on biodiversity conservation and on the resilience of the SES associated to OMYK. Using an integrative dynamical computational model, we tested the hypothesis that: (1) traditional milpa agriculture is compatible with biodiversity conservation in OMYK, and (2) that a balanced diversification between traditional and alternative productive activities is a mechanism that promotes resilience.

## 2 METHODS

### 2.1 Study site

OMYK is located in the northeastern region of the Yucatan Peninsula, Mexico, and it has an extension of 5367 ha. Mean annual temperature in the region is 24.3°C and the mean annual precipitation is 1,120.2 mm (SMN, 2019). The region has a tropical wet and dry climate (Aw2) with a dry season from December to April and a wet season from May to November (CONANP, 2006).

Dominant vegetation is medium semi-evergreen forest in different successional stages (García-Frapolli et al., 2007). The forest forms a heterogeneous landscape composed of a mosaic of vegetation in different successional stages as a result of many years of traditional milpa agriculture and multiple natural disturbances such as hurricanes and forest fires (Bonilla-Moheno, 2008; Rangel-Rivera, 2017).

The site has a high faunistic diversity and is inhabited by multiple endangered species such as spider monkey (*Ateles geoffroyi*), jaguar (*Panthera onca*), puma (*Puma concolor*), howler monkey (*Alouatta pigra*) and others (CONANP, 2006). The spider monkey population inhabiting the reserve has been widely studied (Ramos-Fernández et al., 2018) and has been the main tourist attraction of the site.

The management plan of the reserve formally recognises three user communities: Punta Laguna, Campamento Hidalgo and Nuevo Yodzonot (CONANP, 2006). On this project we focus on Punta Laguna, which given its size and closeness to the reserve has experienced important changes after the decree of the NPA. This community is located in the southeast of the reserve over the roadway Cobá-Nuevo Xcan. Punta Laguna has 136 inhabitants distributed in 28 households. All the inhabitants are indigenous Yucatec Maya (Rivera-Núñez, 2014).

Before the management plan of the reserve entered into force in 2006, households used to manage their resources following a MUS. They used to manage five different land units (milpa, homegardens, secondary forest, old-growth forest and aquatic systems) and implement 13 different productive activities (milpa agriculture, beekeeping, charcoal production, gather firewood, gather medicinal plants, gather wood for home construction, hunting, fishing, ecotourism, home gardening, sheep herding and scientific research assistance; García-Frapolli et al. (2007)). Nowadays, there is a tendency towards an abandonment of the MUS and of the traditional activities, and a growing trend towards a specialization on tourism-related activities (Rios-Beltrán, 2016; García-Frapolli et al., 2012).

There has been continuous research on the site since 1996. The research lines are diverse and include: behavioral ecology of the spider monkeys (Ramos-Fernández et al., 2018), forest succession and restoration (Bonilla-Moheno, 2008; Bonilla-Moheno and Holl, 2010; Bonilla-Moheno, 2010), local systems of nature management (García-Frapolli, 2006; García-Frapolli et al., 2008; Rios-Beltrán, 2016), land cover change analysis (García-Frapolli et al., 2007; Rangel-Rivera, 2017), conservation (Bonilla-Moheno and García-Frapolli, 2012; García-Frapolli, 2015; Rivera-Núñez, 2014) and local institution analysis (García-Frapolli et al., 2013; Rivera-Núñez, 2014).

### 2.2 Description of the model

The model was built integrating an ABM with a BNM. As the BNM can be understood as part of the ABM (Table S1; Figure 1) we describe the whole model following the ODD (Overview, Design concepts and Details) protocol for ABMs (Grimm et al., 2010).

**Figure 1.**
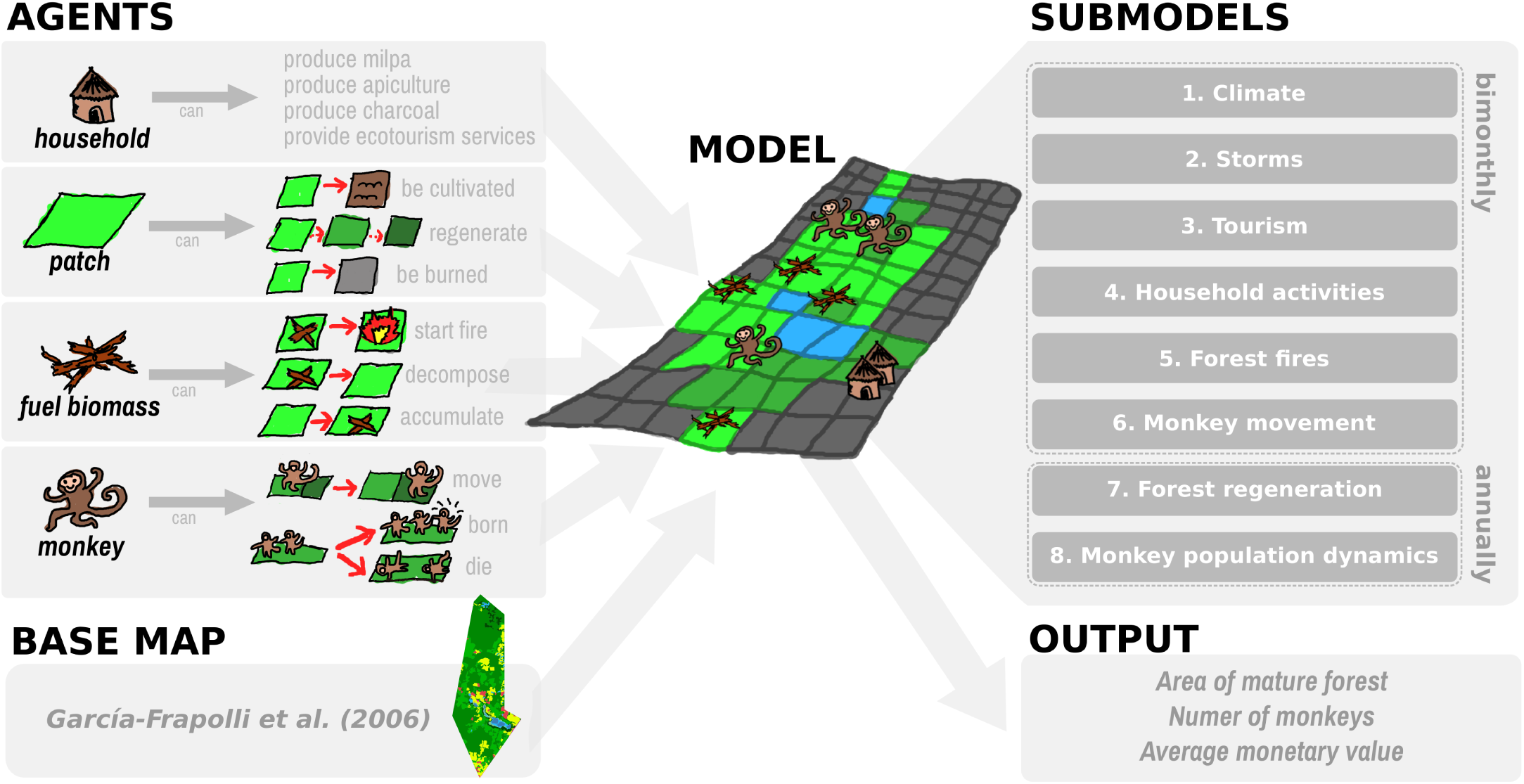
Representation of the model. The model is composed of four different types of agents. The world where the agents interact is based on a land use and vegetation map. The model simulates eight different processes, or submodels, six of which occur every two months (each time step) and the last two only occur annually (each six time steps). The state of the modeled system is monitored through three output variables.

#### 2.2.1 Purpose

The purpose of the model is to explore how different productive and management strategies: (1) affect the capacity of the reserve to conserve biodiversity, defined as the forest area and the size of the spider monkey population, and (2) affect the resilience of some elements of this SES to some frequent disturbances in the region. The model has been designed for scientists and managers, mainly those interested in conservation and forest management in the region, and it is aimed at exploring some hypotheses that have been previously proposed, integrating and synthesizing some of the information that has been generated after more than twenty years of multidisciplinary research in the area.

#### 2.2.2 Entities, state variables and scales

The model consists of four agents: landscape patch, household, monkey and fuel biomass.

Landscape patches represent a square of 3 ha that can be of one of 5 different types: forest, agriculture (milpa), burned, water or household. Patches of type forest, milpa and burn have a successional age that increases year by year, simulating the regeneration of the forest. When the milpa and burned patches reach a successional age of two years they change their type to forest. Patches of type forest and milpa can accumulate fuel biomass, which makes them susceptible to become burned. When a patch is burned or is converted to milpa, its successional age becomes zero and the fuel biomass accumulated on it is consumed. Patches of type forest can be converted into milpas by households that practice agriculture. After a milpa has been used for two consecutive years, it is abandoned and allowed to regenerate, and the household opens a new plot (García-Frapolli et al., 2007). Water-type patches represent water bodies and the household-type patches represent family homes (built-up constructions). Water and household-type patches remain fixed throughout the simulation.

Households have a fixed position in the virtual ABM world. They can only work on four different productive activities: milpa agriculture, apiculture, charcoal production and provide ecotourism services. The activities that households perform remain fixed throughout the simulation. Households obtain a monetary value from each of the productive activities they do. Disturbances, such as hurricanes, forest fires and fluctuations in tourism, have an effect on the productive activities that households do, thus affecting their monetary value.

A monkey agent represents a spider monkey. These can live and move in the forest-type patches with a successional age greater than or equal to 30 years. Monkeys can be born or die following a local logistic growth equation (see 2.2.7.8. Monkey population dynamics).

Fuel biomass represents an aggregate of decomposing biomass that can ignite a fire. Fuel biomass accumulates on patches after storms and hurricanes. Each fuel biomass unit has a duration time before it disappears. While the fuel biomass is present on a patch, it can cause it to burn.

One time step of the model represents two months (see 7.1. Climate). The model is run on a 37 x 94 grid where each grid cell represents 3 ha of landscape.

#### 2.2.3 Process overview and scheduling

Within a bimester, a sequence of processes takes place in the following order. First, the weather conditions of the bimester are determined (i.e., rainy season or dry season). Second, depending on the weather conditions, different meteorological events may occur (e.g., tropical storms, hurricanes) and depending on the type of event a proportion of patches accumulates fuel biomass. Third, the flow level of tourists visiting the reserve is determined. Fourth, households make their productive activities and calculate how much monetary value they obtain. Fifth, forest fires can occur on patches with fuel biomass accumulated. Finally, monkeys can move from patch. Annually (i.e., every 6 iterations), two additional processes follow: patches regenerate and spider monkey population dynamics take place.

As simulations are performed along discrete time steps, all the state variables of the agents are updated synchronously.

#### 2.2.4 Design concepts

##### 2.2.4.1 Emergence

The interaction and feedback between the social and ecological systems emerge as a result of land use changes and the productive activities done by the households.

##### 2.2.4.2 Adaptation/Objectives/Learning/Prediction/Sensing

Agents do not present any adaptive behavior, objectives, learning, prediction nor sensing. The set of activities a household does makes up a household strategy, but households do not change their strategy along the simulation.

##### 2.2.4.3 Interactions

Households interact indirectly with monkeys through the fragmentation and habitat loss caused by milpa agriculture. Monkeys indirectly affect the monetary values of the households by regulating the tourist flow. Households also interact directly with the patches by cultivating them and by removing fuel biomass. Spider monkeys are also affected by high levels of tourist flow.

##### 2.2.4.4 Stochasticity

The occurrence of the different meteorological events is stochastically determined based on occurrence probabilities (see sections below). The selection of the patches that accumulate fuel biomass is random. Tourist flow has a stochastic component. The selection of new parcels for cultivation, the site for placing the beehives and the fuel biomass used by the households is partly random. Forest fires are modeled using a stochastic percolation model. And the movement of the monkeys is partly random.

##### 2.2.4.5 Collectives

Contiguous patches of the same type and within the same range of successional age aggregate into patch neighborhoods. Monkeys living in the same patch neighborhood are aggregated into a local population with its own population dynamics.

##### 2.2.4.6 Observation

The average annual monetary value of the productive activities done by the households (Table S3) is observed to assess the economic state of the households. The mature forest area (successional age >50 years), the vegetation type that harbours the greater species richness and most rare species (Bonilla-Moheno, 2008), and the total number of spider monkeys, are observed to assess the capacity of the reserve to conserve biodiversity.

#### 2.2.5 Initialization and input

According to literature reports (Rivera-Núñez, 2014), the number of households is set to 28 and the number of monkeys is set to the maximum possible in each habitable patch neighborhood. To create the virtual world, we used a 2003 land use and vegetation map (García-Frapolli et al., 2007). This map was rasterized and inserted into the landscape grid. The successional age of the patches is set to the minimum age according to the range to which they belong on the base map (2-7, 8-15, 16-29, 30-50, >50 years). The model only considers the area that is occupied by the reserve, so all the grid cells outside of the polygon of the reserve are ignored. The model parameters were set to the values shown in Table S2.

#### 2.2.6 Input

The model does not use input data to represent time-varying processes.

#### 2.2.7 Submodels

##### 2.2.7.1 Climate

Weather conditions are modeled using BNM. This submodel is based on the submodel of the same name of López-Martínez (2017) and is used to give a temporality to the model. The submodel is composed of three nodes: temperature, pressure and precipitation. The interpretation of the state of the nodes, regulation functions and their explanations are shown in Table S1. The dynamics of this submodel generates a periodic attractor of length 6 where the node precipitation remains in one same state for three consecutive states of the attractor and in a qualitatively different state during the next three states. This dynamics is interpreted as an annual climatic dynamic with 6 months of rainy season and 6 months of dry season. This climatic dynamic is characteristic of the region, so each state of the attractor is mapped to one bimester of the year. This mapping allowed us to locate different events throughout the year.

##### 2.2.7.2 Storms

Tropical storms and hurricanes are a frequent disturbance in the region. The strong winds of hurricanes generate defoliation, snapping and uprooting of trees (Bonilla-Moheno, 2010). So, after these disturbances there is a high accumulation of fire biomass that can generate forest fires in the subsequent dry seasons. To simulate this, a proportion of affected patches (*P* (*s*)) is calculated as a function of the speed of the wind (*s*). We suppose that the relation between these two variables follows a sigmoid function:

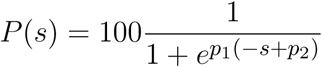

where *p*_1_ and *p*_2_ are parameters that determine the shape and position of the inflection point, respectively. The speed of wind (*s*) takes a value depending on the type of event that occurs on a bimester: no storm (10 mph), tropical storm (20 mph) and hurricanes from category one to five (74, 96, 111, 130 and 157 mph). On the rainy season storms and hurricanes can take place with certain occurrence probability estimated with data from NOAA (2019) (Table S2). When an event occurs a unit of fuel biomass is created on an amount of patches of type milpa or forest randomly selected equivalent to the proportion of affected patches (*P* (*s*)). When created, fuel biomass units are assigned a time of duration from normal distribution (mean duration of fuel biomass, Table S2).

##### 2.2.7.3 Tourism

The tourist flow to the reserve is modeled using BNM. This submodel is composed of two nodes, tourism and tourismH, that together represent three levels of tourists flow: no tourists, low season flow and high season flow. The interpretation of the state of the nodes, regulation functions and their explanation are shown in Table S1. With these rules we obtain a dynamic with two bimester of high season and four of low season that is similar to the tourist dynamic in the region.

As hurricanes generally reduce the flow of tourists (García-Frapolli et al., 2012), we assumed that if a hurricane occurs on a bimester, then no tourists arrive (i.e., the nodes tourism and tourismH turn off). Likewise, as the main tourist attraction at the site are spider monkeys (García-Frapolli et al., 2013), we assumed that if there are not at least 20 monkeys at a ratio of 3 km from the households location then no tourists arrive.

Finally, a stochastic factor was added to simulate the effect of economic and social disturbances that also reduce the number of tourists. This was simulated through a parameter that with certain probability avoids a high flow of tourists (i.e., the node tourismH is kept turned off; probability of high flow of tourists, Table S2).

##### 2.2.7.4 Household activities

Household activities are also modeled using BNM. This submodel is composed of seven nodes: openMilpa, plantMilpa, youngMilpa, adultMilpa, harvestMilpa, harvestApiculture and charcoalProduction. The interpretation of the state of the nodes, regulation functions and their explanation are shown in Table S1. With these rules we obtained a dynamic that allows to locate the realization of the households’ productive activities throughout the year (see 3.1. Submodels build with boolean network modelling). In contrast to the other submodels built with BNM, whose nodes are global (i.e., one set of nodes for all the system), the nodes that constitute this submodel are unique for each household considered in the simulation (i.e., each household has its own set of nodes).

To spatially simulate the production of milpa agriculture, when a household opens a milpa it chooses a plot equivalent to 3 ha within a radio of 3 km from is location. Milpas are only opened in patches with successional age greater than or equal to 5 and less than 50 years.

To simulate the effect of the hurricanes on the agriculture it is supposed that if a hurricane occurs and the milpa plot of a household is affected by it (i.e., accumulates fossil fuel), then the plants and harvests are lost (i.e., youngMilpa, adultMilpa and harvestMilpa nodes turn off). Likewise, to simulate the effect of hurricanes on apiculture, it is supposed that a household places their beehives in a patch in a radius of 3 km from its location, and if a hurricane occurs and this patch is affected, then the beehives are damaged and there will be no harvests during the next two years. Finally, it is also supposed that households that produce charcoal consume one unit of fuel biomass in a radius of 3 km from its location.

Each bimester, households calculate their total bimonthly monetary value from the values shown in Table S3.

##### 2.2.7.5 Forest fires

This submodel uses a modified version of the percolation model of the model library of NetLogo 6.0.4 (Wilensky, 2006). In the model forest fires can only occur with certain occurrence probability during the dry season estimated with data form (CONABIO, 2019) (Table S2). If a forest fire occurs, then one of the patches with most fuel biomass accumulated ignites, and with certain probability (burning probability of patches, Table S2) the four neighbour patches containing fuel biomass can also ignite. Then, each burned patch repeats the process.

##### 2.2.7.6 Monkey movement

Spider monkeys can move with certain probability that depends on the successional age of the patch they are on (probability of permanence, Table S2). If a monkey moves, then it randomly shifts to a neighbouring patch with successional age greater than or equal to 30 years. If there are monkeys on patches of type milpa or burned, then they move to one of their nearest patches with successional age greater than or equal to 30 years.

##### 2.2.7.7 Forest regeneration

For simplification purposes, we supposed that the regeneration of the forest is deterministic and that the only factor affecting the successional state of the patches is time. So, each year the patches of type milpa, burn and forest increase their successional age.

##### 2.2.7.8 Monkey population dynamics

This submodel is based on the submodel named “Animal meta-population dynamics” of Iwamura et al. (2014). First, patch neighborhoods (set of neighbor patches within the same range of successional age) of late successional forest (30-50 years) and mature forest (>50 years) are formed. Then, for each of these neighborhoods the local population size (*N*_*t*_) is estimated using the logistic growth equation:

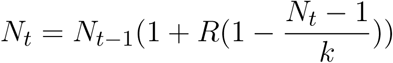

Where *N*_*t*−1_ is the local population size in the time *t* −1, *R* is the discrete intrinsic growth rate and *k* is the carrying capacity of the neighborhood. *R* was obtained from the literature, and *k* for the nth neighborhood, denoted *k*_*n*_, is calculated in the next way:

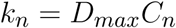

Where *D*_*max*_ is the maximum density of a patch and *C*_*n*_ is the number of patches that form the nth neighborhood. *D*_*max*_ values were obtained from the literature (Table S2).

Finally, as it has been proposed that a high flow of tourists can have an effect on the monkey population, (for instance, the noise generated by tourists and the opening of new trails may distress the monkeys affecting their breeding areas and feeding habits; García-Frapolli et al. (2007)), we supposed that the discrete intrinsic growth rate (*R*) diminishes by a sixth for each bimester that there is a high flow of tourists.

### 2.3 Implementation

The model was implemented on NetLogo 6.0.4. The code can be found on: https://github.com/laparcela/OMYKmodel

### 2.4 Calibration

We made a manual calibration of the following four parameters related to the extent of the forest fires for which no information was founded on the literature:

1. burning probability of patches (values explored: 0.4, 0.5, 0.56, 0.6, 0.7),
2. sigmoid function parameter 1 (*p*_1_) (values explored: 0.05, 0.045, 0.040, 0.035, 0.03, 0.025, 0.02),
3. sigmoid function parameter 2 (*p*_2_) (values explored: 74, 96, 111), and
4. mean duration of fuel biomass (values explored: 6, 9, 12).

We explored a total of 315 different treatments that represent all the possible combinations of the selected values for each parameter (Figure S1). We run each treatment 100 times over a time span of 12 years (from 2003 to 2015) in which households do not produce milpa agriculture (as it was prohibited in 2006). We selected the combination of values for the parameters by applying the following criteria:

1. that the total burned area registered by the model was similar to the total area burned observed by Rangel-Rivera (2017) form 2003 to 2015 (i.e., that the empirical data was inside the interquartile range of the model results).
2. that the dispersion of the model results was from the ten smallest.
3. that the burning probability of patches was the least possible. This was done with the objective of avoiding the fires that affect the complete reserve that appear in longer simulations.
4. that the percentage of affected patches when no storm occurs was the lowest possible. This was done to avoid the Forest Fire and Storms submodels to work independently from each other.

After applying this criteria hierarchically (i.e., first applying criteria 1, next criteria 2, and so on) we chose the values for the parameters that are shown in Table S2.

### 2.5 Sensitivity analysis

We carried out a sensitivity analysis on six parameters as a way of examining the consistency (verification) and the general behaviour of the model. The explored variables and the explored values for each were:

1. patch size: 1 and 3 ha;
2. burning probability of patches: from 0 to 1, by increments of 0.1;
3. sigmoid function parameter 1 (*p*_1_): from 0.01 to 0.05, by increments of 0.005;
4. sigmoid function parameter 2 (*p*_2_): 74, 96, 111 and 130;
5. mean duration of fuel biomass: 6, 9, 12 and 15 bimesters;
6. standard deviation of duration of fuel biomass: 1, 3, 6, 9 and 12 bimesters;
7. probability of high flow of tourists: from 0 to 1, by increments of 0.1;
8. minimum number of monkeys for tourism: from 10 to 35, by increments of 5.

We followed the “one at a time” method, in which one parameter is varied at a time while keeping the other fixed on the values shown in Table S2. We ran 30 simulations for each treatment during a virtual time span of 200 years (so that the output variables stabilize) in which households produce the four productive activities. We assessed the sensitivity visually looking for noticable changes in the model results.

### 2.6 Validation test

Model results were compared with empirical data of the different vegetation and land use classes in 2015 (Rangel-Rivera, 2017) and estimated monkey population size of 2015 (Spaan, 2017). We ran 100 simulations over a virtual time span of 12 years (from 2003 to 2015) in which households do not produce milpa agriculture. Model parameters were fixed on the values shown in Table S2. The values of the four parameters for which no data was found in the literature and no calibration was made, were chosen arbitrarily. We look for values that would make simulations faster and that would not generate noticeable changes in the output variables according to the results of the sensitivity analysis.

### 2.7 Scenarios

To assess the effect of different management strategies on the capacity of the reserve to conserve biodiversity we explore the 16 different possible combinations of the four productive activities that can be implemented by the households (we only explored the cases where all the households implemented the same set of activities). Each treatment was run 30 times during a time span of 50 years. Combinations were classified and averaged following to the characterization of strategies of García-Frapolli et al. (2007) (Table S4):

1. Traditional: combinations where households produce agriculture but no ecotourism.
2. Mixed: combinations where households produce both milpa and ecotourism.
3. Service Oriented: combinations where households dedicated to ecotourism but no agriculture.
4. Other: combinations that did not correspond to any of the previous ones.

To explore the effect of different management strategies on the resilience of the SES to some frequent disturbances we ran the next set of scenarios:

1. We increased the frequency of hurricanes and tropical storms by arbitrarily multiplying the probabilities of occurrence of tropical storms and of each type of hurricane by 3.
2. We increased the frequency of forest fires by arbitrarily multiplying their probability by 3.
3. We reduced the flow of tourism by arbitrarily reducing the probability of high flow to 0.3.

Each scenario was run 30 times during a time span of 50 years.

## 3 RESULTS

### 3.1 Submodels built with boolean network modelling

The sequence of system states (periodic attractor) recovered by the three submodels built with BNM (just considering a single household) is shown in Figure 2. Each of the states of the attractor can be interpreted as a “photography” of the approximate conditions of the SES in a given bimester. For instance, the second state of the attractor may be interpreted as the March-April bimester, a bimester of the dry season (node precipitation turned off) in which a high flow of tourists can arrive (nodes tourism and tourismH turned on); in this bimester a household generally prepares the plot for the milpa (node openMilpa turned on); and also in these months a household can obtain good harvests of apiculture and produce charcoal (nodes harvestApiculture and charcoalProduction turned on). Similarly, the fifth state of the attractor may be interpreted as a rainy season bimester (September-October), with a low flow of tourists on which the milpa is almost ready to be harvested and on which households can produce charcoal.

**Figure 2.**
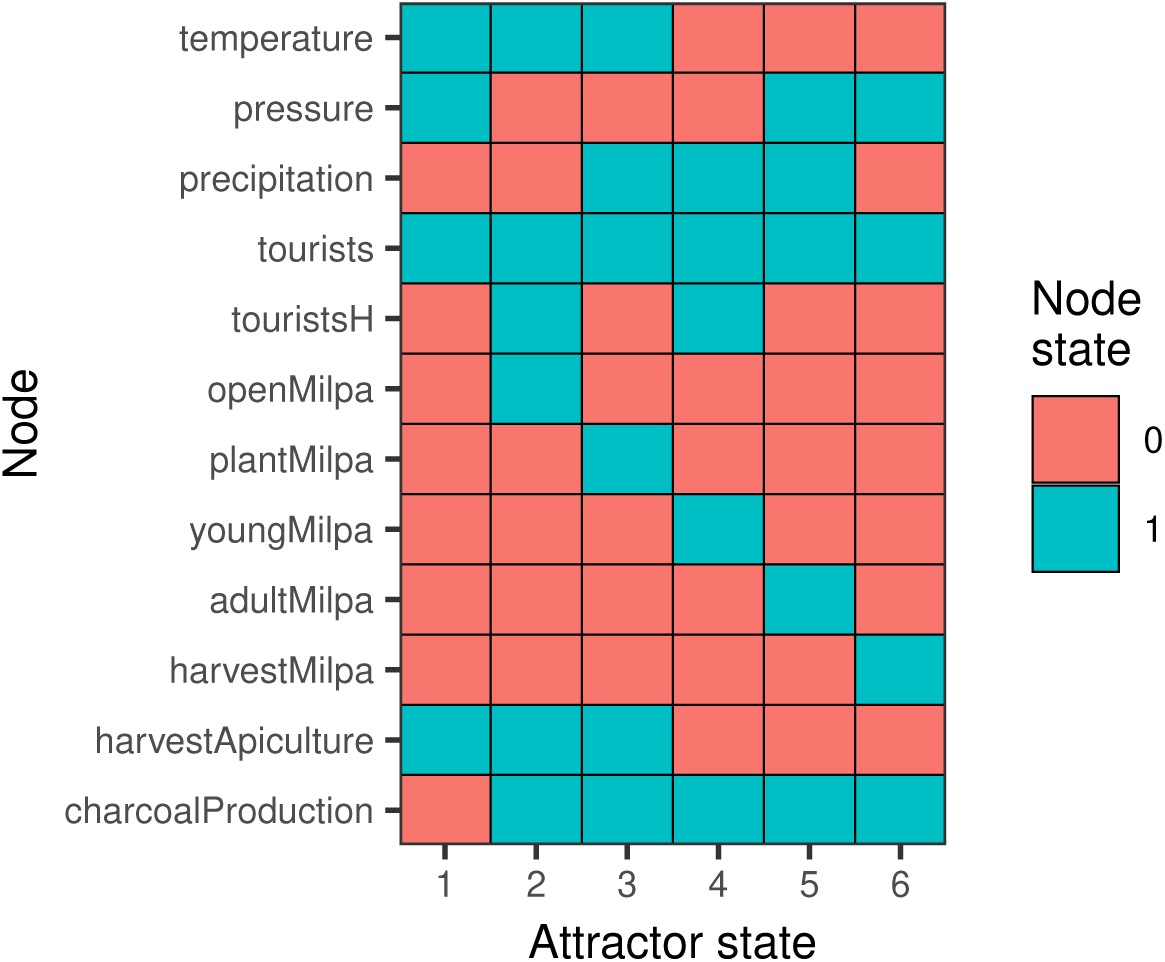
Attractor recovered by the submodels built with boolean network modelling. Rows represent each node of the network, columns are the attractor states and each color represents a different state of the node. Each of the attractor states of this cyclic attractor of period six can be interpreted as a snapshot of the approximate state of the SES in a given bimester (See Subsection 3.1. Submodels built with boolean network modelling).

### 3.2 Sensitivity analysis

We observed the following behavior patterns for the three output variables:

1. Mature forest area. In most simulations this variable presents three events of remarkable growth as a result of the initial conditions of the mode and of the wide length of the age categories used to classify the forest. These events take place at 21, 35 and 50 years and correspond to the moment when the patches that started at 30, 16 and 2 years, respectively, reach the mature forest category. After these three events the variable stabilizes.
2. Number of monkeys. In most cases this variable decreases slightly during the first 21 years. Next, it gradually increases until it stabilizes.
3. Average monetary value. In most simulations this variable stabilizes since the beginning and fluctuates around certain equilibrium values. In some cases the equilibrium values gradually decrease until they reach a new, lower equilibrium.

Most output variables were sensible to the modification of the explored parameters. Mature forest area and the total number of monkeys were sensible to the burning probability of patches (Figure 3), the mean duration of the fuel biomass (Figure 4), the two parameters of the sigmoid function (Figures S2-S3), and patch size (Figure S4). Average monetary value was sensible to burning probability of patches (Figure 3), patch size (Figure S4), the probability of high flow of tourists (Figure S5) and the minimum number of monkeys for tourism (Figure S6).

**Figure 3.**
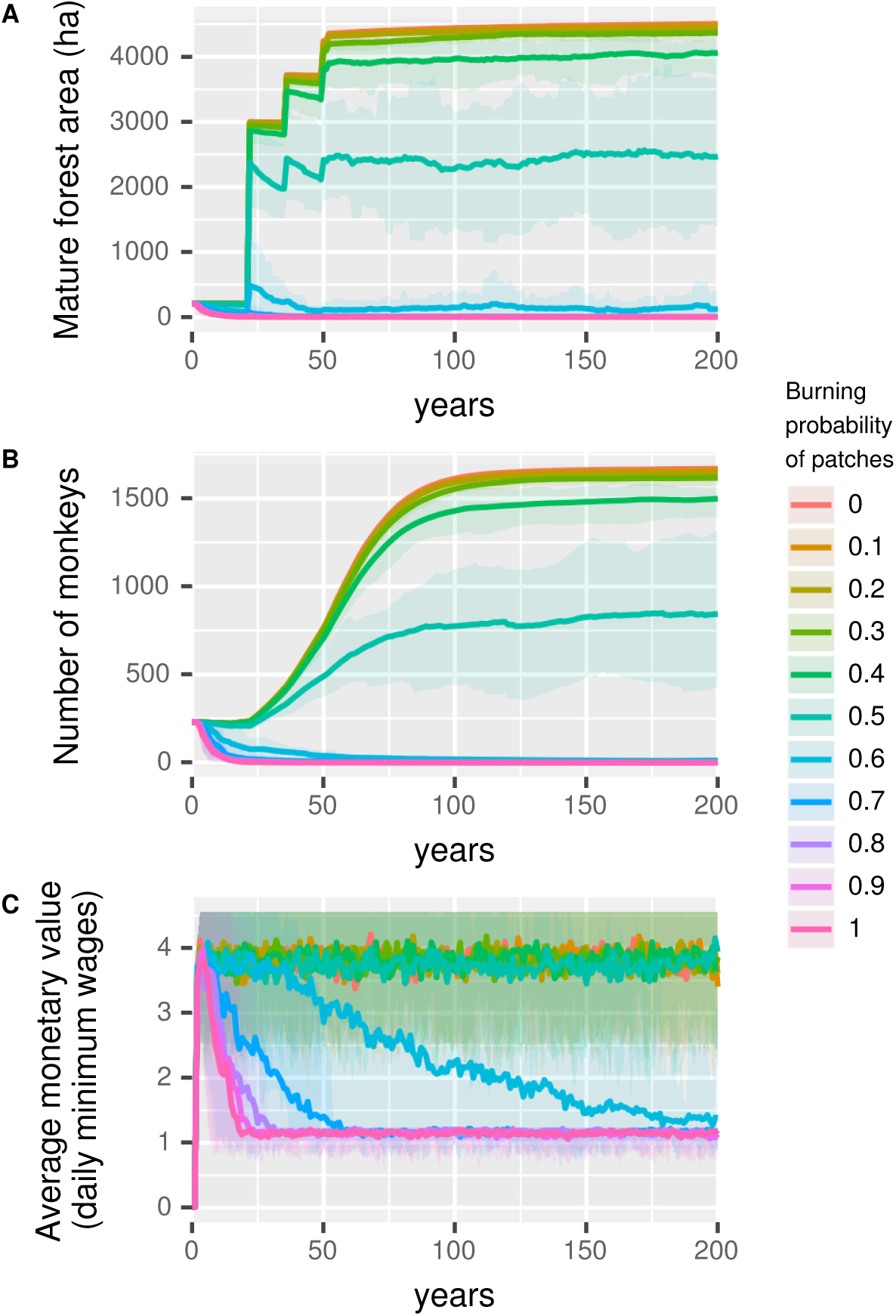
Sensitivity analysis of the burning probability of patches. Each panel represents the change in an output variable for a 200 year period and 30 simulation runs for each value of the burning probability of patches: A): Mature forest area in ha (forest greater than 50 years of successional age). B): total number of monkeys. C): average monetary value of activities done by the households in daily minimum wages. Lines represent the mean and upper and lower shadows represent 5th and 95th quartiles, respectively, for each value of the burning probability of patches (different colors).

**Figure 4.**
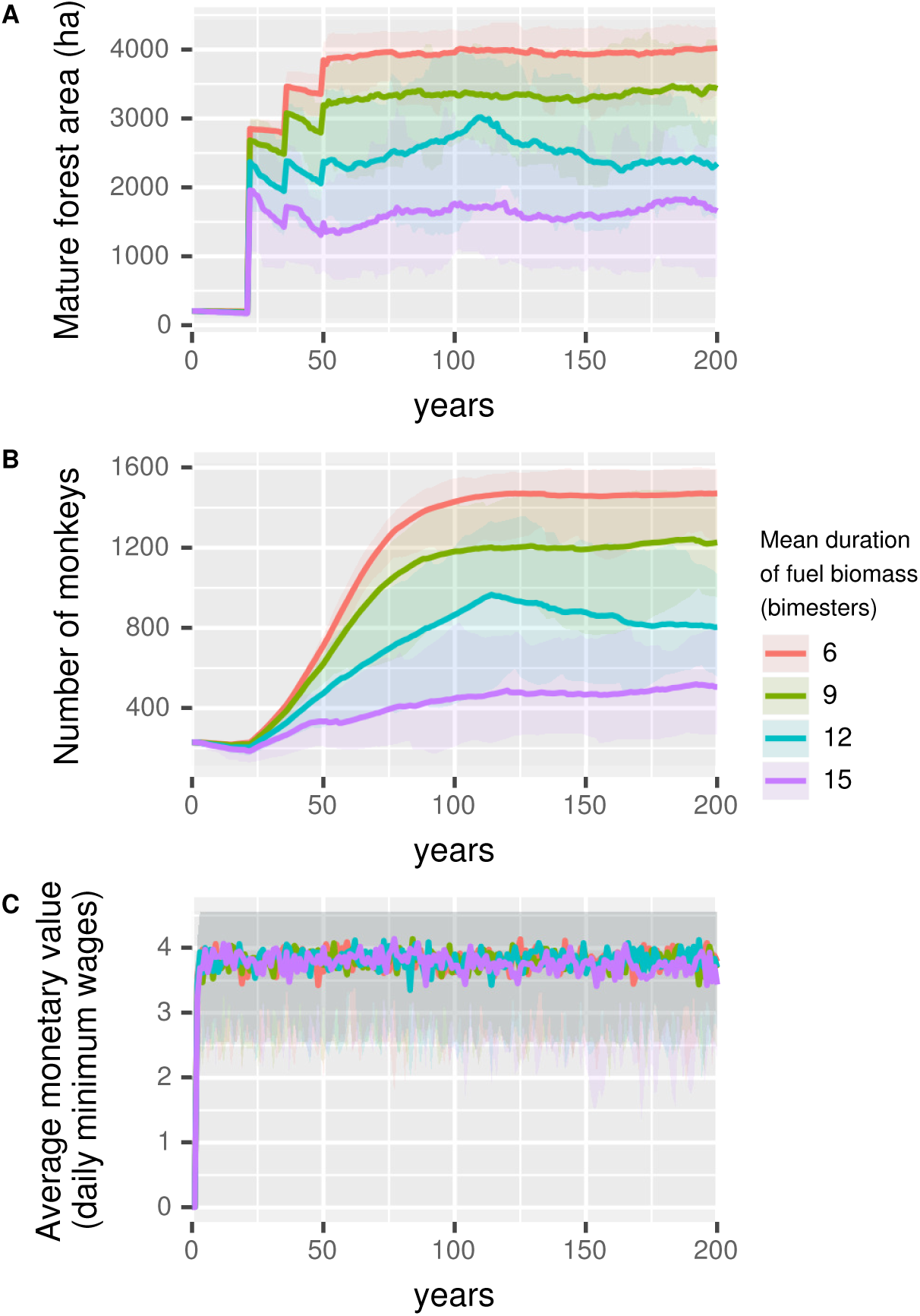
Sensitivity analysis of the mean duration of fuel biomass (bimesters). Each panel represents the change in an output variable for a 200 year period and 30 simulation runs for each value of the mean duration of fuell biomass. A): Mature forest area in ha (forest with successional age greater than 50 years). B): total number of monkeys. C): average monetary value of activities done by the households in daily minimum wages. Lines represent the mean and upper and lower shadows represent 5th and 95th quartiles, respectively, for each value of the mean duration of fuel biomass (different colors).

### 3.3 Validation test

The model reproduced relatively well the area of forest of 2-15, 16-29 and 30-50 years (Figure 5). Nevertheless, the model overestimated the area of mature forest and the total number of monkeys.

**Figure 5.**
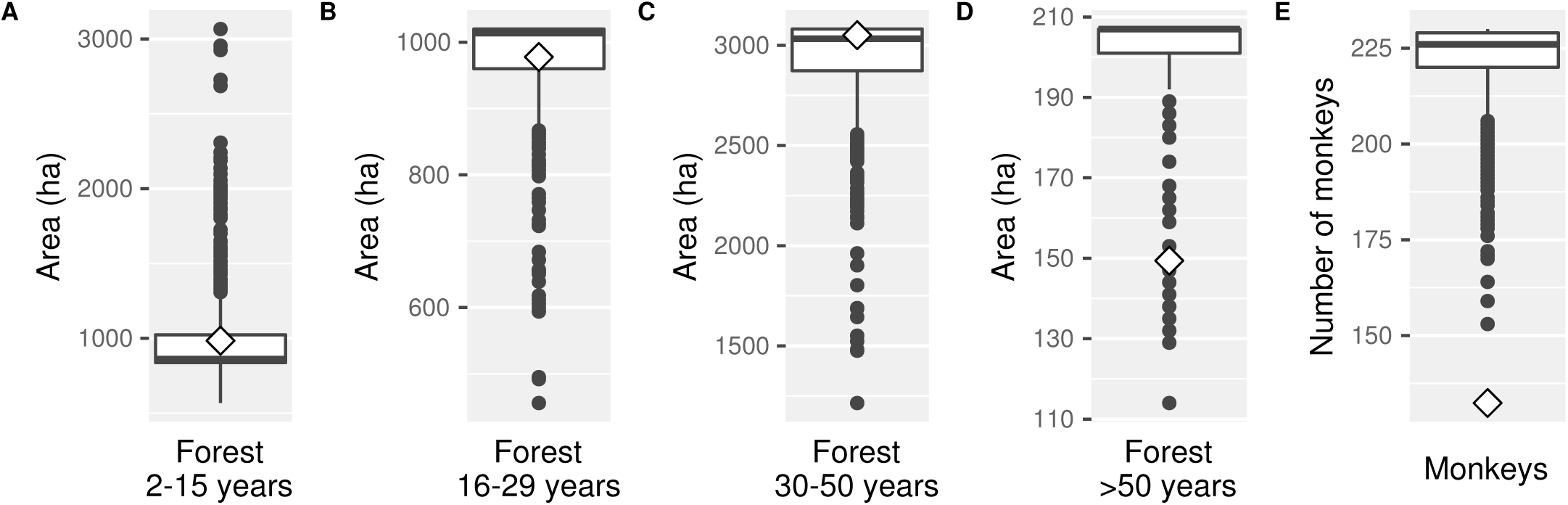
Validation test results. Results of simulations and empirical values for five different variables. A): area of forest with successional age between 2 and 7 years. B): area of forest with successional age between 16 and 29 years. C): area of forest with successional age between 30 and 50 years. D): area of forest with successional age greater than 50 years. E): total number of monkeys in the reserve. Boxplots are the model outputs after 12 years (from 2003 to 2015) for 100 simulation runs. Diamonds represent the empirical observed values for different forest types in 2015 and the total estimated number of monkeys in 2015.

### 3.4 Scenarios

In the scenario without any extra disturbance (Figure 6 first column), mature forest area was greater for Service Oriented and Other strategies. However, there was a growth trend of this variable in all four strategies. Similarly, there was a growth trend of the monkey population in all the strategies, with a larger population increase in Other strategy, followed by the Service Oriented and Traditional strategies, and finally by the Mixed strategy. Average monetary value was markedly higher for the strategies including ecotourism, though the variability was higher as the households depended more on this activity (Table S5).

**Figure 6.**
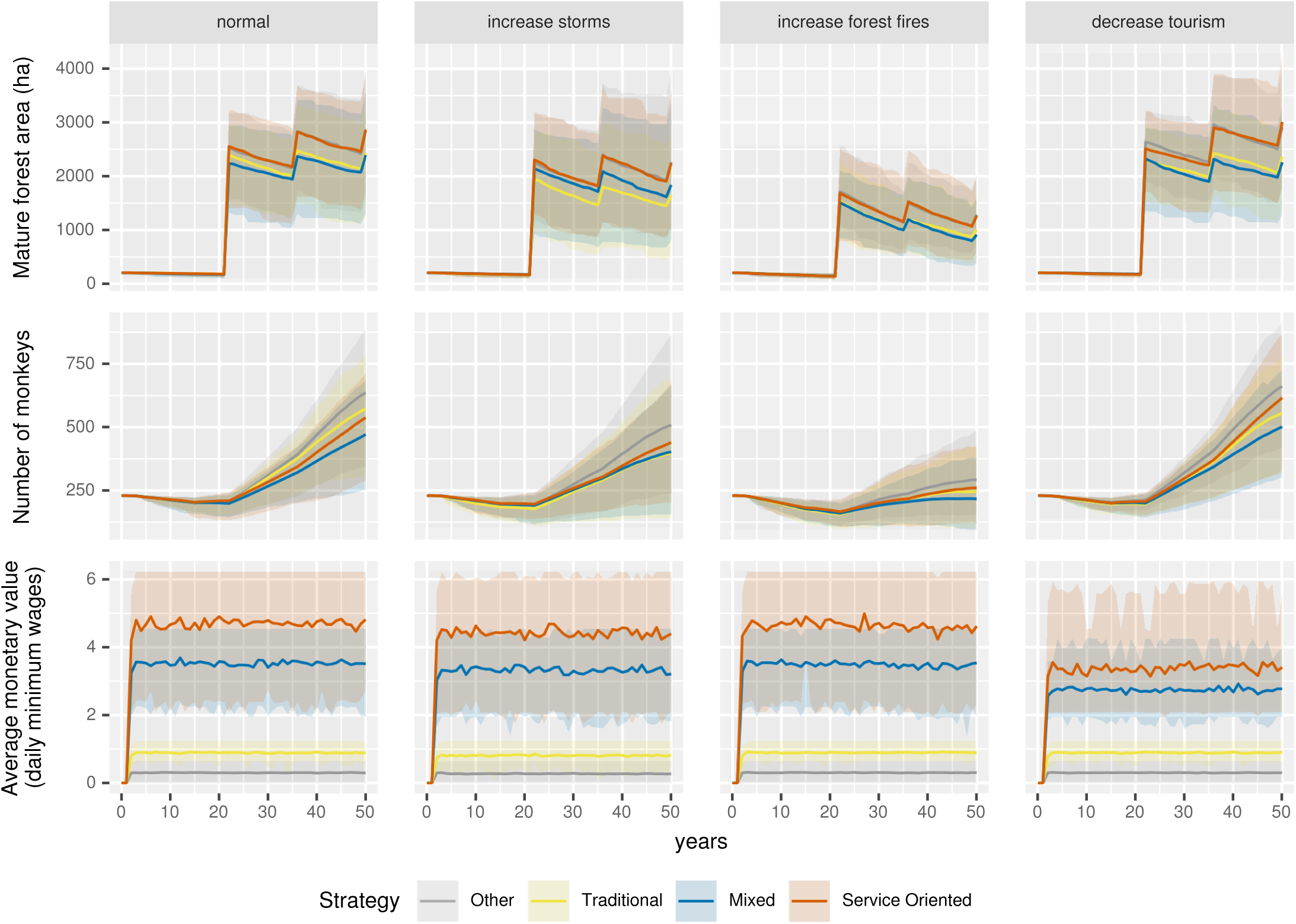
Model results for different strategies under different scenarios. Each column of graphs represents a different scenario and each row a different output variable. The four scenarios shown are: a scenario with normal level of all disturbances, a scenario with an increase in the storms and hurricane occurrence probability, a scenario with an increase in the probability of occurrence of forest fires and a scenario with a decrease in the probability of the flow of tourists. First row: Mature forest area in ha (forest with successional age greater than 50 years). Second row: total number of monkeys. Third row: average monetary value of activities done by the households in daily minimum wages. Lines represent the mean and upper and lower shadows represent 5th and 95th quartiles, respectively, for different strategies (in colors), each ran over a 50 year period and for 30 simulation runs.

When the storm occurrence probability was increased there was just a slight reduction of the trajectories of the three output variables for all strategies (Figure 6 second column). In the scenario of increased forest fires (Figure 6 third column) there was a considerable reduction of mature forest area and monkey population in all strategies, but there were just slight changes in the average monetary value. In contrast, when tourist flow was reduced (Figure 6 fourth column), there were no noticeable changes in the trajectories for the mature forest area and the number of monkeys, yet there was a remarkable reduction of the average monetary value for the strategies that included this activity. The reduction of the aggregated monetary value under all disturbances was higher for the Service Oriented strategy than for the Mixed strategy (Figure 7, Table S5).

**Figure 7.**
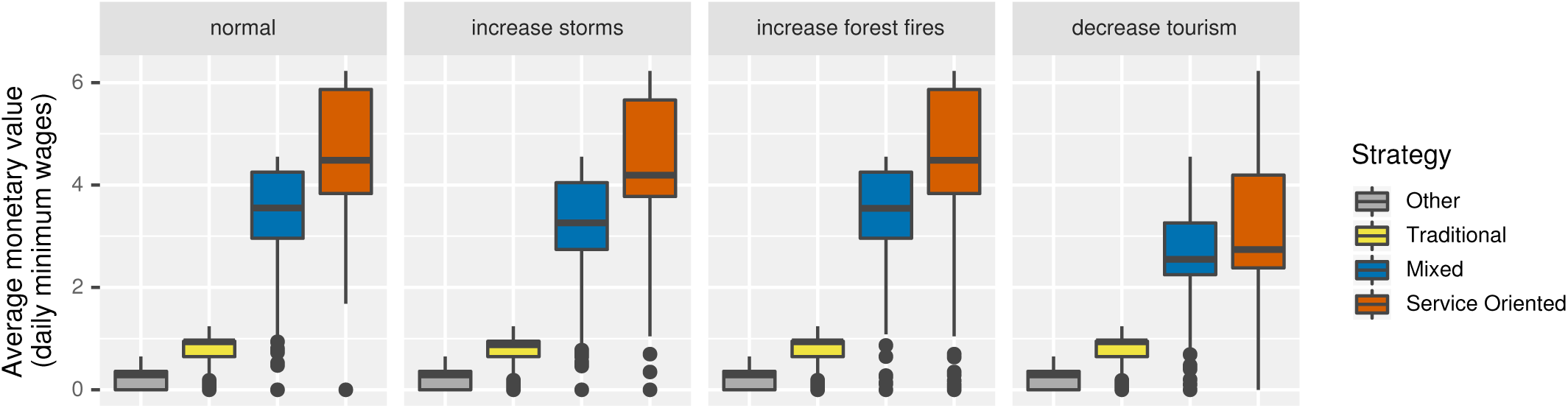
Aggregate average monetary value for different strategies under different disturbance scenarios. The four scenarios shown are: a scenario with normal level of all disturbances, a scenario with an increase in the storms and hurricane occurrence probability, a scenario with an increase in the probability of occurrence of forest fires and a scenario with a decrease in the probability of the flow of tourists. Boxplots represent the yearly average monetary value of different strategies (in colors) for 30 simulations each ran over a 50 year period.

## 4 DISCUSSION

The model proposed here enabled the formal integration of ecological and social data and information collected over the past 20 years, and allowed us to test how different management strategies affect the biodiversity conservation and socio-ecosystemic resilience in the long term. We show that traditional activities like swidden milpa agriculture are compatible with forest and biodiversity conservation, and that a balance of traditional and alternative activities is a way to promote resilience.

The results of the sensitivity analysis suggest that the model has an important degree of uncertainty associated with the modification of its parameters. Nevertheless, the behaviors observed in the output variables can be explained by the assumptions of the conceptual model (i.e., the mechanisms proposed in subsection 2.2.7). This fact increases the confidence on the model implementation. In the case of the validation test, the model was able to reproduce quite well the empirically observed area of successional forest categories but failed to reproduce the observed mature forest area and estimated number of monkeys. This may be partly due to the unusually large forest fires that took place between 2006 and 2011, which damaged large areas at the north of the reserve affecting particularly one of the main patches of mature forest (Rangel-Rivera, 2017). It is important to notice that the calibration and the validation test were done with data of the same study. This can explain the good fit of some of the model results. To better test the predictive capacity of the model it will be necessary to compare it with future independent studies. Even if the model could not be correctly validated and even if it may not be useful to make precise quantitative predictions, it reproduces fairly well the mid-term qualitative system behavior described in the literature. As such, the model proved to be a useful tool to explore the joint effect of the considered variables and gain new insights of this SES.

The small effect of milpa agriculture on the prevalence and increment of mature forest and monkey population suggests that the production of traditional milpa agriculture is compatible with the conservation of biodiversity in OMYK. These results agree with model predictions obtained by García-Frapolli et al. (2007). These authors, using a probabilistic model, found no significant differences in the reduction of mature forest in scenarios with and without milpa agriculture in OMYK. This compatibility of milpa agriculture and biodiversity conservation is due to multiple factors. First, the low rate at which milpa agriculture was done prior to its prohibition in 2006 did not compromised the extension of remaining older successional forest. Second, before the decree of the reserve, the local community, incentivized by ecotourism, had already agreed on not using the mature forest surrounding the lagoon for agricultural activities (García-Frapolli et al., 2007). This initiative allowed the conservation of one of the main patches of mature forest used by the spider monkeys. As the model results suggest, this zonification, that did not eliminate completely milpa agriculture from the reserve, may have been a sufficient measure to conserve the spider monkey population, as well as many other species that share their habitat. And third, the spider monkeys may be more flexible to the degradation of their habitat than previously thought (Spaan, 2017). In the reserve, spider monkeys are not restricted to mature forest as they also use and travel on late successional forest fragments (30-50 years old; Ramos-Fernández and Ayala-Orozco (2003); Ramos-Fernandez et al. (2013)). This allows them a greater mobility to exploit a wide range of environments in a fragmented habitat. More generally, in spite of the general perception that agricultural activities are a threat to primate biodiversity, there is evidence of primate presence on a diversity of agroecosystems (Estrada et al., 2006).

As our results show, the incorporation of ecotourism has brought significant economic benefits to the households. Regular and relatively high incomes from this activity have significantly reduced the need of the household members to search for temporary work outside their communities, improving family well-being (García-Frapolli et al., 2007). However, the abandonment of traditional activities and the MUS reduces the resilience of this SES. The greater resistance to all disturbances and the lesser variability of the Mixed strategy, support the hypothesis that a balanced diversified strategy between traditional and alternative productive activities is a mechanism that promotes resilience. This is explained by the greater functional redundancy and response diversity that this strategy promotes (Biggs et al., 2012). For instance, while in a Service Oriented strategy a reduction of the tourist flow may be strongly perceived by the household, in a Mixed strategy the income reduction can be buffered by the goods obtained on other activities. In general, these results agree with studies that suggest that economic diversification helps rural households reduce poverty and vulnerability (Ellis, 2008; Martin and Lorenzen, 2016; Thulstrup, 2015) and studies that document the greater vulnerability of tourism specialization (Tao and Wall, 2009; Su et al., 2016).

Further, specialization also threatens food sovereignty and biocultural diversity. As specialization on ecotourism increases, there is a greater dependency of the communities on external agents and a loss of control over their subsistence. This means, for example, that communities are forced to buy products that they previously produced and decided how to produce. The abandonment of milpa agriculture is also worrying, given that it is a central component of the Yucatec Maya identity and, as such, it has associated a great diversity of knowledge, practices and beliefs which may become lost (Barrera-Bassols and Toledo, 2005; Lyver et al., 2019). Similarly, tourism dynamics have led to a process of acculturation, as shown by the way that the traditional ceremonies and rituals have been commodified and are now commercial products for the tourists (García-Frapolli et al., 2018).

The disturbance that most affected the mature forest and the monkey population was the increase in forest fires. The importance of this disturbance on the SES was also observed in the sensitivity analysis when modifying the burning probability of patches and the mean duration of biomass fuel. When varying the burning probability of patches we obtained the behaviour characteristic of the percolation models, with a threshold that when exceeded, prevents the growth of the mature forest and the monkey population. It is worth noting that in order to reproduce the empirically observed total burned area, in the calibration, it was found that this parameter has to be set near the critical point. Indeed, it has been proposed that some ecological systems organize near the critical points, particularly regarding the occurrence of forest fires (Ricotta et al., 1999). Similarly, when varying the mean duration of fuel biomass, we found a clear reduction of the mature forest area and of the spider monkey population. This was due to the greater extent of the forest fires given the greater proportion of affected patches at a given point of time. This result suggests that some traditional activities that involve the reduction of the volume of fuel material, as the gathering of firewood and wood for construction and the controlled burning for traditional agriculture, may be important mechanisms to prevent large-scale forest fires. However, as some of these activities are mainly seen as detrimental to the environment and some evidently involve fire risk, they have been prohibited in the reserve (CONANP, 2006).

Throughout most of the 20th century, environmental policy on fire on most countries, including Mexico, was based on a fire suppression approach (Martínez-Torres et al., 2016). This approach is responsible for the widely negative perception on fire use on traditional agriculture, as it blamed the practices of rural and indigenous people as the main cause of forest fires (Martínez-Torres et al., 2016). Currently, Mexico environmental policy on fire is on a transition towards an integrated management approach which recognises the role that fires could play in ecosystems (Gutiérrez Navarro et al., 2017). However, as environmental policy on fire responds mainly to ecological concerns, it continues to ignore the practices and knowledge on fire use of rural and indigenous people (Gutiérrez Navarro et al., 2017; Martínez-Torres et al., 2016; Monzón-Alvarado et al., 2014). In the case of the Yucatec Mayas, “wind tenders” (*yum ik’ob* in Yucatec Maya) are the people who carry out the burns in the milpa production. Wind tenders have developed a variety of techniques to control fire, preventing it to expand to the surrounding fields by placing firebreaks, while also assuring the complete conversion of biomass into ashes (Nigh and Diemont, 2013). Given the need of effective large-scale fire prevention strategies and the profound traditional ecological knowledge of the Maya people, it seems essential to revalue local practices and knowledge regarding fire management. We do not suggest that traditional or local practices are ideal, but that they should not be ignored and that the local communities’ perspectives, needs and knowledge should not be excluded from fire management plans. In general, our findings suggest that forest fire prevention strategies in NPAs need to go beyond the recognition and prohibition of productive activities with possible detrimental effects on the environment, and also recognise the role that these activities may play in ecosystem management and fire prevention.

The model presented in this work is a simplified representation of a complex SES and has multiple limitations. We discuss four of these. Firstly, the model only considers the area occupied by the reserve. This implies that it ignores all the ecological and social processes that take place outside its administrative borders. For instance, although the habitat of the monkeys may be recovering within the reserve, other processes (e.g., intensification of agricultural activities or expansion of urban areas) may be occuring outside of it that may jeopardize the long-term permanence of monkey populations (e.g., loss of connectivity between habitats in the region). Secondly, the model does not consider the human demographic changes. Thus, we could not explore how population growth may promote the expansion of agriculture and urban areas and the overall consequences of such expansion. However, population growth in OMYK does not necessarily imply an expansion of the agricultural land given the existence of alternative productive activities (García-Frapolli et al., 2007). Thirdly, the income distribution is rarely equitable and accessible to all, as assumed in the model. In OMYK, while the community of Punta Laguna has highly benefited from tourism, other communities have been excluded or have not consolidated initiatives that allow them to enjoy the same benefits (Rivera-Núñez, 2014). Even among the members of Punta Laguna, there has been an exclusion of the households that have remained more attached to their traditional activities (García-Frapolli et al., 2018). Fourth, the model does not consider some of the main current forces of change in this SES. Since the reserve’s decree, inhabitants have faced strong conflicts over land tenure and usufruct. There is a conflict between the community members of Punta Laguna and some members of the ejido assembly. For years the ejido, a collective form of land tenure, has demanded a greater economic gratification of ecotourism activities, while the local community members have refused to do so (García-Frapolli et al., 2013). This conflict has led to eviction threats to the local community members. Similarly, in the past years there was an illegal attempt to change the land tenure regime, from collective to private, of the lands occupied by the reserve. With these changes, the land surrounding the reserve would be modified form collective to private parcels, which could then be sold (García-Frapolli et al., 2007). This would enable the entry of external agentes, quite likely from the tourism development and real estate industries.

To our knowledge, this is the first work on the study of SESs that explicitly integrates BNMs and ABMs. One of the main constraints of the latter is that they require large amounts of empirical data to be parameterized (Iwamura et al., 2014). In contrast, BNMs require no or few parameters, making them a suitable tool for modeling scarcely characterized systems (Saadatpour and Albert, 2013). In this sense, the qualitative approach offered by the BNMs can facilitate and simplify the construction of models for the study of SES. Our model can be used as a guide or support tool for future integrative studies in OMYK. Similarly, the model can be used as a guide towards the development of support tools in participatory processes. This type of tools can help in the communication of knowledge, promote collective reflection and learning, and thus facilitate collective and adaptive decision-making for building sustainable management practices and effective governance (Braasch et al., 2018; D’Aquino et al., 2003).

In conclusion, the results of this study suggest that: (1) conservation strategies that do not exclude traditional productive activities can be compatible with biodiversity conservation, and (2) a balanced economic diversification between traditional and alternative activities, is a strategy that promotes the economic resilience of rural households. In addition, the model presented here shows how computational modeling and the SES perspective are effective means of integrating and synthesizing information from different sources. Finally, our work also illustrates some of the main limitations of systems perspective and computational modelling on the study of SES, such as the difficulty in considering political, cultural and historical features.

## AUTHOR CONTRIBUTIONS

LGGJ, MB and GRF conducted the research. EGF, MBM and CRR provided data and maps for the model. LGGJ wrote original draft. All authors reviewed and edited the paper.

## ACKNOWLEDGMENTS

We thank the community of Punta Laguna for their support of academic research over the years. We also thank Celene Espadas, Fernanda Figueroa, Fernanda Ríos-Beltrán, Tlacaelel Rivera-Núñez and Ernesto Vega for their commentaries on previous versions of this work. M.B. acknowledges financial support from UNAM-DGAPA-PAPPIIT (IN207819). G.R.F. acknowledges support from CONACYT (grant 157353) and Instituto Politécnico Nacional.

