## Supplementary Material for "Multiple resource use strategies confer resilience to the socio-ecosystem in a protected area in the Yucatan Peninsula, Mexico"

**Table S1.** Interpretation, transition function and explanation of nodes composing the submodels built with boolean networks.

| Submodel | Node | Inerpretation of node state | Transition function | Explanation |
|---|---|---|---|---|
| Climate | *temperature* ($tmp$) | 1: high mean temperature<br>0: low mean temperature | $tmp_{t+1} = \neg pcp_t$ | The absence of rainy clouds generates an increase in air temperature. Likewise, rainy clouds favor solar radiation reflection generating a cooling in the earth's surface (López-Martínez, 2017). |
| | *pressure* ($prs$) | 1: high atmospheric pressure<br>0: low atmospheric pressure | $prs_{t+1} = \neg tmp_t$ | When air temperature increases, it goes up in the air column reducing atmospheric pressure. Likewise, with lower temperatures the air goes down increasing the atmospheric pressure (López-Martínez, 2017). |
| | *precipitation* ($pcp$) | 1: high average precipitation<br>0: low average precipitation | $pcp_{t+1} = \neg prs_t$ | Low atmospheric pressure promotes the entrance of humid winds from the ocean that may form rainy clouds. Likewise, high atmospheric pressure is associated with dry seasons (López-Martínez, 2017). |
| Tourism | *tourism* ($tur$) | 1: low flow of tourists<br>0: no tourists | $tur_{t+1} = pcp_t \vee \neg pcp_t$ | A low flow of tourists can arrive all year round. |
| | *tourismH* ($tuh$) | 1:high flow of tourists<br>0: no high flow of tourists | $tuh_{t+1} = tur_t \wedge tmp_t \wedge (prs_t \vee pcp_t)$ | Tourism in the region has a strong temporality (SECTUR, 2019). We fixed the high seasons in the second (May-April) and fourth (July-August) bimesters. |
| Household activites | *openMilpa* ($omi$) | 1: household opens milpa<br>0: household doesn't open milpa | $omi_{t+1} = tmp_t \wedge \neg prs_t \wedge \neg pcp_t$ | The milpas are burned prior to the rainy season, between March and April (Emerson, 1953). |
| | *plantMilpa* ($pmi$) | 1: household plants milpa<br>0: household doesn't plant milpa | $pmi_{t+1} = omi_t$ | The milpa is planted at the beginning of the rainy season, after the burning of the milpa (Emerson, 1953). |
| | *youngMilpa* ($ymi$) | 1: milpa has young crops<br>0: milpa doesn't have young crops | $ymi_{t+1} = pmi_t \wedge pcp_t$ | Milpa harvest can start in August and end until Febraruy (Emerson, 1953). We supposed that after six months of growth the milpa is harvested and the only factor necessary for the growth of the plants is the rain water. |
| | *adultMilpa* ($ami$) | 1: milpa has adult crops<br>0: milpa doesn't have adult crops | $ami_{t+1} = ymi_t \wedge pcp_t$ | |
| | *harvestMilpa* ($hmi$) | 1: household harvests milpa<br>0: household doesn't harvest milpa | $hmi_{t+1} = ami_t \wedge pcp_t$ | |
| | *harvestApiculture* ($hap$) | 1: household harvest beehives<br>0: household doesn't harvest beehives<br>0: household doesn't harvest beehives | $hap_{t+1} = \neg pcp_t$ | The best harvests of honey are obtained during the dry season when the principal melliferous species flower (Rico-Gray and Chemas, 1991). |
| | *charcoalProduction* ($chp$) | 1: household produces charcoal<br>0: household doesn't produce charcoal | $chp_{t+1} = \neg hmi_t$ | The charcoal production is mainly done in the free time of other activities (García-Frapolli, 2006). We supposed that the milpa harvest, a very demanding activity, impedes the production of charcoal. |

**Table S2.** Values and sources of the main parameters of the model.

| Submodel | Parameter | Value | Source |
|---|---|---|---|
| Household activities | Number of households | 28 | Rivera-Núñez (2014) |
| | Time of usage of a same milpa plot | 2 años | García-Frapolli et al. (2007) |
| | Maximum ratio of activities | 3 km | García-Frapolli et al. (2007) |
| | Minimum fallow period | 5 años | García-Frapolli et al. (2008) |
| | Milpa plot size | 3 ha | García-Frapolli et al. (2007) |
| Forest fires | Patch burning probability | 0.5 | Estimated in Calibration / Sensitivity analysis |
| | Probability of forest fire occurrence in dry season | 0.093 | Estimated from CONABIO (2019)[a] |
| | Patch size | 3 ha | Sensitivity analysis |
| Storms | Probability of occurrence tropical storm | 0.083 | Estimated from NOAA (2019)[b] |
| | Probability of occurrence hurricane category 1 | 0.010 | Estimated from NOAA (2019)[b] |
| | Probability of occurrence hurricane category 2 | 0.007 | Estimated from NOAA (2019)[b] |
| | Probability of occurrence hurricane category 3 | 0.010 | Estimated from NOAA (2019)[b] |
| | Probability of occurrence hurricane category 4 | 0.007 | Estimated from NOAA (2019)[b] |
| | Probability of occurrence hurricane category 5 | 0.000 | Estimated from NOAA (2019)[b] |
| | Sigmoid function parameter 1 ($p_1$) | 0.025 | Estimated in Calibration / Sensitivity analysis |
| | Sigmoid function parameter 2 ($p_2$) | 74 | Estimated in Calibration / Sensitivity analysis |
| | Mean duration of fuel biomass | 12 bimesters | Estimated in Calibration / Sensitivity analysis |
| | Standard deviation of duration of fuel biomass | 3 bimesters | Sensitivity analysis |
| Tourists | Probabilidy of high flow of tourists | 0.7 | Sensitivity analysis |
| | Minimum number of monkeys for tourists to arrive | 15 monkeys | Sensitivity analysis |
| Monkey movement | Probability of staying in late successional forest (30-50 years) | 0.19 | Estimated from Spaan (2017) |
| | Probability of staying in mature forest (>50 years) | 0.81 | Estimated from Spaan (2017) |
| Monkey population dynamics | Discrete intrinsic growth rate ($R$) | 0.10 | Robinson and Redford (1986); Ross (1988) |
| | Maximum density of monkeys in late successional forest | 5.70 $\frac{ind}{km^2}$ | Spaan (2017) |
| | Maximum density of monkeys in mature forest | 38.20 $\frac{ind}{km^2}$ | Spaan (2017) |

[a] Estimated from a total of 24 records from the period 2000-2017.

[b] To calculate these probabilities a total of 36 records of storms and hurricanes were obtained after applying the following filters: (1) that it was a record from 1916 to 2016, (2) that the record, with a diameter buffer of 100 km intersected the polygon of the reserve, and (3) that it was the record with the maximum wind speed of a same storm or hurricane.

**Table S3.** Bimonthly monetary value per activity. Estimated from data of García-Frapolli (2006) and Rivera-Núñez (2014). We assumed that households that provide ecotourism services but don't produce milpa are specialized in ecotourism activities and consequently earn more from this activity.

| Productive activity | Monetary value (daily minimum wages) |
|---|---|
| Milpa harvest (i.e., node *harvestMilpa* turned on) | 3.89 |
| Apiculture harvest (i.e., node *harvestApiculture* turned on) | 0.73 |
| Charcoal production (i.e., node *charcoalProduction* turned on) | 0.36 |
| Low season (i.e., node *tourism* turned on and *tourismH* turned off) for household producing milpa and providing ecotourism services | 1.31 |
| Low season for household providing ecotourism services and not producing milpa | 2.09 |
| High season (i.e., nodes *tourism* and *tourismH* turned on) for household producing milpa and providing ecotourism services | 7.32 |
| High season for household providing ecotourism services and not producing milpa | 12.55 |

**Table S4.** Explored scenarios and their classification.

| Combination of activities | Strategy |
|---|---|
| - - - - | Other |
| - - - T | Service oriented |
| - - C - | Other |
| - - C T | Service oriented |
| - A - - | Other |
| - A - T | Service oriented |
| - A C - | Other |
| - A C T | Service oriented |
| M - - - | Traditional |
| M - - T | Mixed |
| M - C - | Traditional |
| M - C T | Mixed |
| M A - - | Traditional |
| M A - T | Mixed |
| M A C - | Traditional |
| M A C T | Mixed |

M: milpa agriculture

A: apiculture

C: charcoal production

T: ecotourism

**Table S5.** Mean and mean variation of average monetary value (in daily minimum wages) under different disturbances for different strategies. $\bar{x}$: mean average; $\bar{s}$: mean standard deviation; $\overline{C.V.}$: mean variation coefficient.

| Strategy | normal | | | increase storms | | | increase forest fires | | | decrease tourism | | |
|---|---|---|---|---|---|---|---|---|---|---|---|---|
| | $\bar{x}$ | $\bar{s}$ | $\overline{C.V.}$ | $\bar{x}$ | $\bar{s}$ | $\overline{C.V.}$ | $\bar{x}$ | $\bar{s}$ | $\overline{C.V.}$ | $\bar{x}$ | $\bar{s}$ | $\overline{C.V.}$ |
| Other | 0.30 | 0.22 | 73.93 | 0.27 | 0.21 | 78.36 | 0.30 | 0.22 | 73.68 | 0.29 | 0.22 | 73.81 |
| Traditional | 0.86 | 0.23 | 26.60 | 0.78 | 0.25 | 32.45 | 0.86 | 0.22 | 25.69 | 0.86 | 0.23 | 26.35 |
| Mixed | 3.39 | 0.73 | 21.54 | 3.19 | 0.81 | 25.36 | 3.36 | 0.77 | 22.96 | 2.64 | 0.70 | 26.42 |
| Service Oriented | 4.51 | 1.15 | 25.61 | 4.25 | 1.30 | 30.65 | 4.45 | 1.27 | 28.54 | 3.25 | 1.09 | 33.68 |

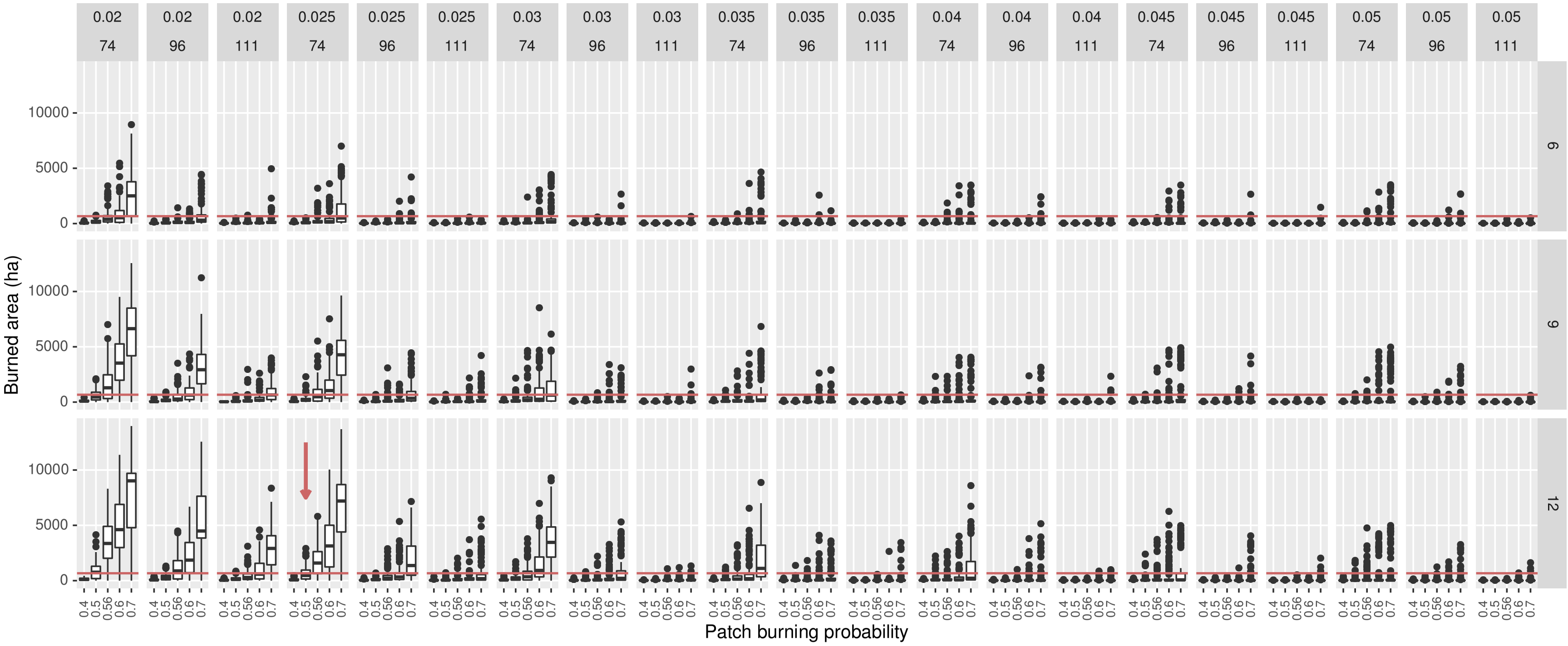


**Figure S1.** Calibration of four parameter with unknown values. Each graph column in the panel represents a combination of values of the two parameters that control the sigmoid function (sigmoid function parameter 1 ($p_1$) and sigmoid function parameter 2 ($p_2$)). Each graph row represent a different value for the mean duration of fuel biomass. Inside each panel, each column represents a value of the patch burning probability. The red line is on 667.2 ha, the total burned area observed between 2003 and 2005 by Rangel-Rivera (2017). This was the value we used as reference for the selection of the values of these parameters. An arrow indicates the combination of parameters that was choosen.

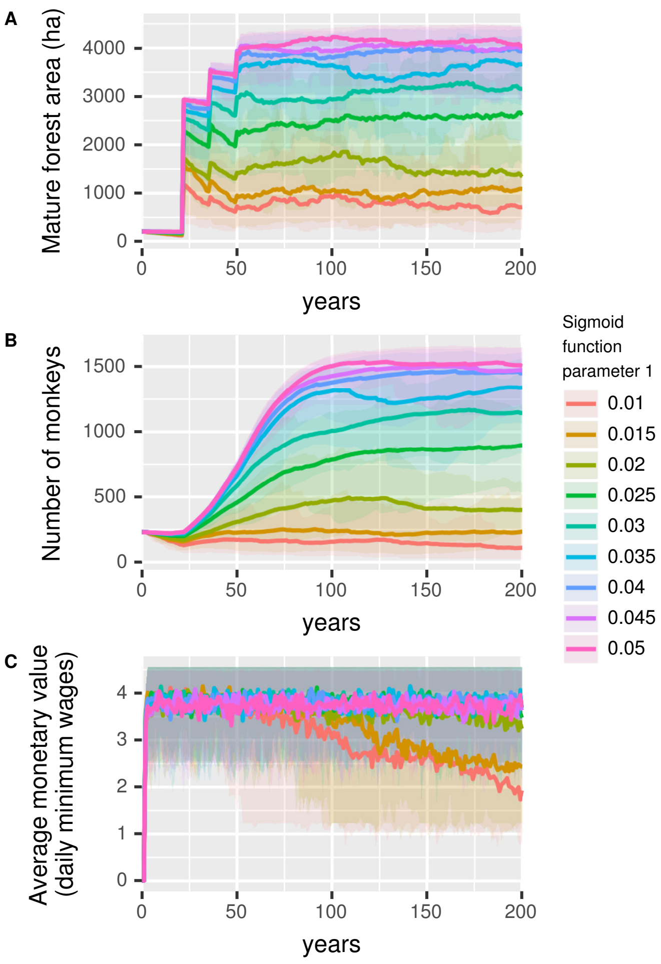


**Figure S2.** Sensitivity analysis of sigmoid function parameter 1 ($p_1$). Each panel represents the change in an output variable for a 200 year period and 30 simulation runs for each value of the sigmoid function parameter 1. A): Mature forest area in ha (forest with successional age greater than 50 years). B): total number of monkeys. C): average monetary value of activities done by the households in daily minimum wages. Lines represent the mean and upper and lower shadows represent 5th and 95th quartiles, respectively, for each value of the sigmoid function parameter 1 (different colors).

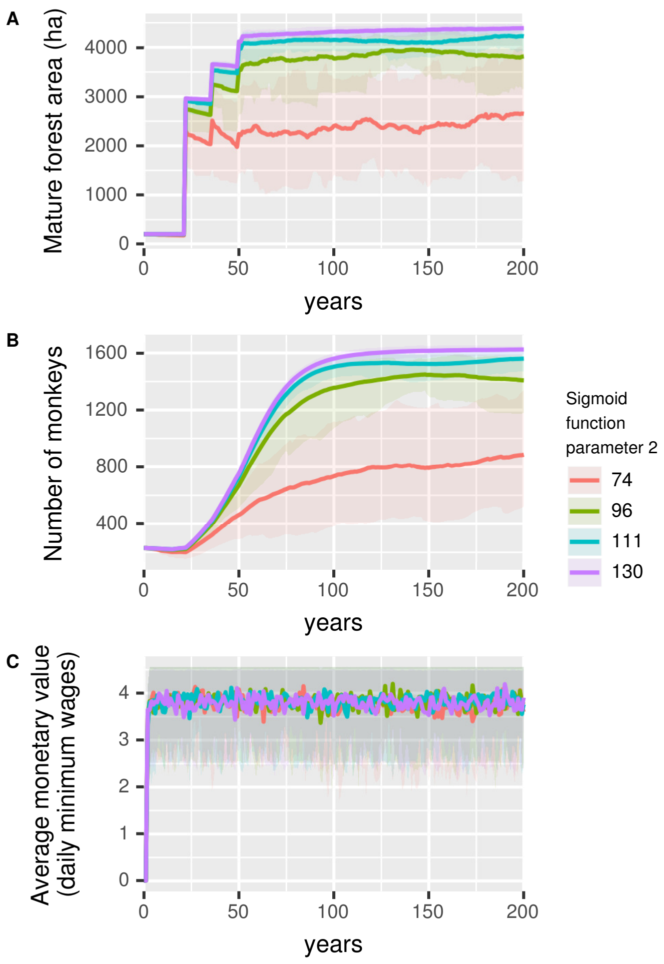


**Figure S3.** Sensitivity analysis of sigmoid function parameter 2 ($p_2$). Each panel represents the change in an output variable for a 200 year period and 30 simulation runs for each value of the sigmoid function parameter 2. A): Mature forest area in ha (forest with successional age greater than 50 years). B): total number of monkeys. C): average monetary value of activities done by the households in daily minimum wages. Lines represent the mean and upper and lower shadows represent 5th and 95th quartiles, respectively, for each value of the sigmoid function parameter 2 (different colors).

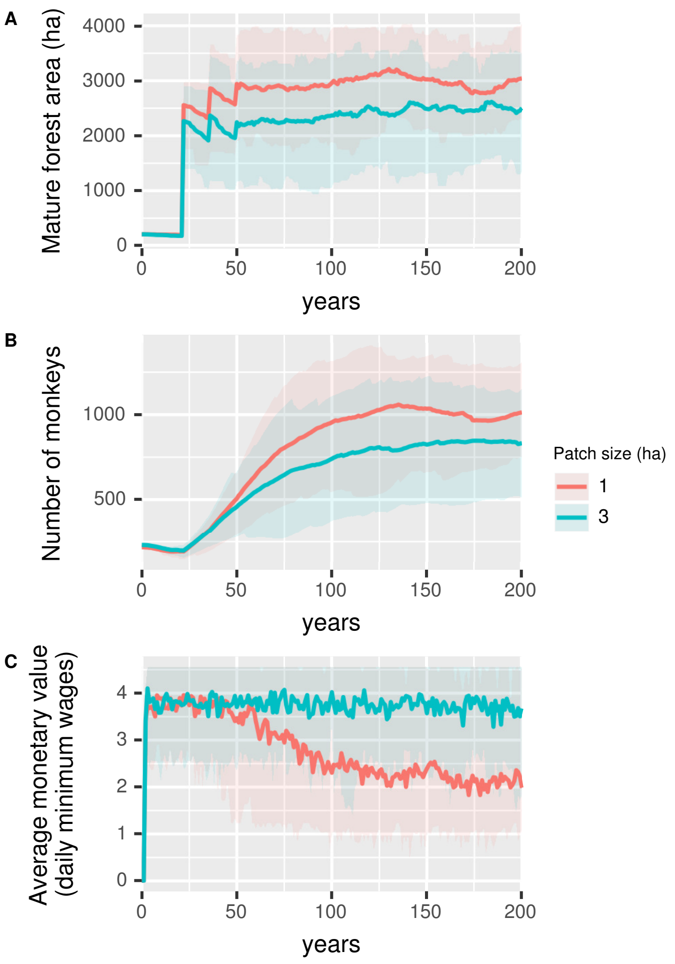


**Figure S4.** Sensitivity analysis of patch size (ha). Each panel represents the change in an output variable for a 200 year period and 30 simulation runs for each value of the patch size. A): Mature forest area in ha (forest with successional age greater than 50 years). B): total number of monkeys. C): average monetary value of activities done by the households in daily minimum wages. Lines represent the mean and upper and lower shadows represent 5th and 95th quartiles, respectively, for each value of the patch size (different colors).

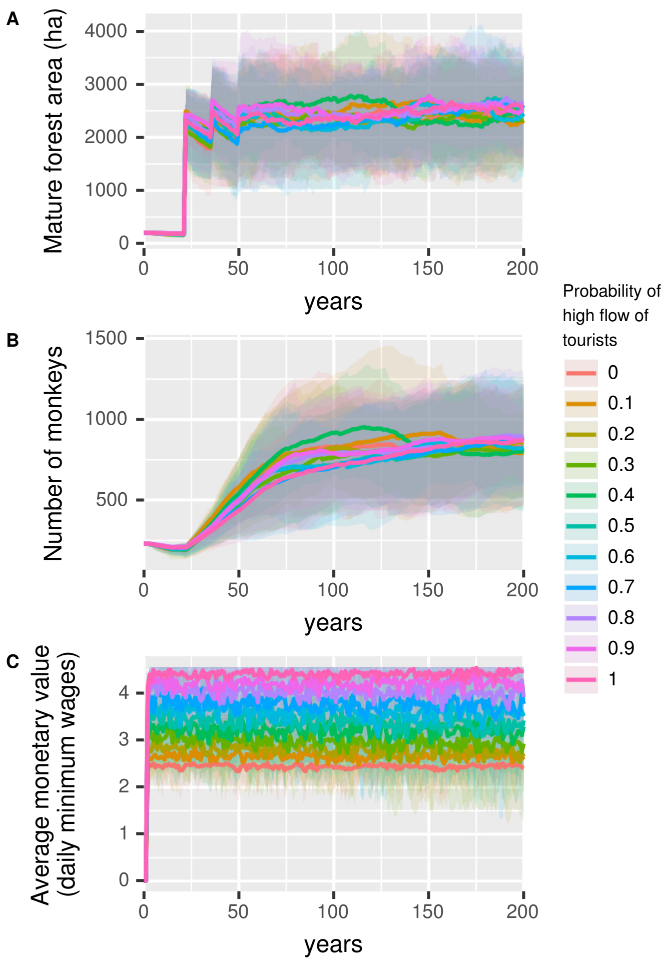


**Figure S5.** Sensitivity analysis of the probability of high flow of tourists. Each panel represents the change in an output variable for a 200 year period and 30 simulation runs for each value of the probability of high flow of tourists. A): Mature forest area in ha (forest with successional age greater than 50 years). B): total number of monkeys. C): average monetary value of activities done by the households in daily minimum wages. Lines represent the mean and upper and lower shadows represent 5th and 95th quartiles, respectively, for each value of probability of high flow of tourists (different colors).

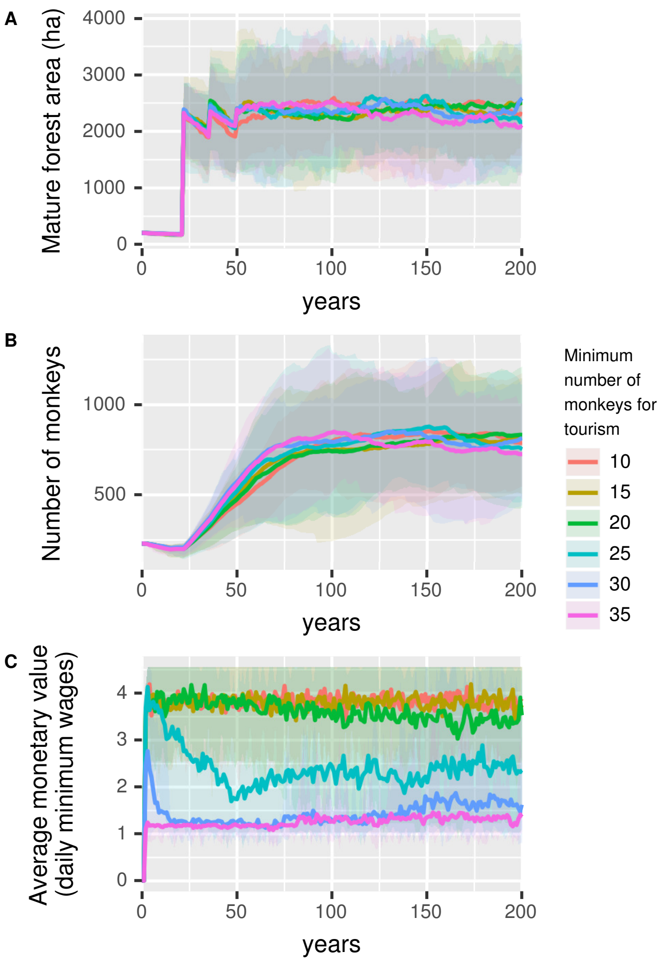


**Figure S6.** Sensitivity analysis of minimum number of monkeys for high flow of tourism. Each panel represents the change in an output variable for a 200 year period and 30 simulation runs for each value of the minimum number of monkeys for high flow of tourism. A): Mature forest area in ha (forest with successional age greater than 50 years). B): total number of monkeys. C): average monetary value of activities done by the households in daily minimum wages. Lines represent the mean and upper and lower shadows represent 5th and 95th quartiles, respectively, for each value of tminimum number of monkeys for high flow of tourism (different colors).

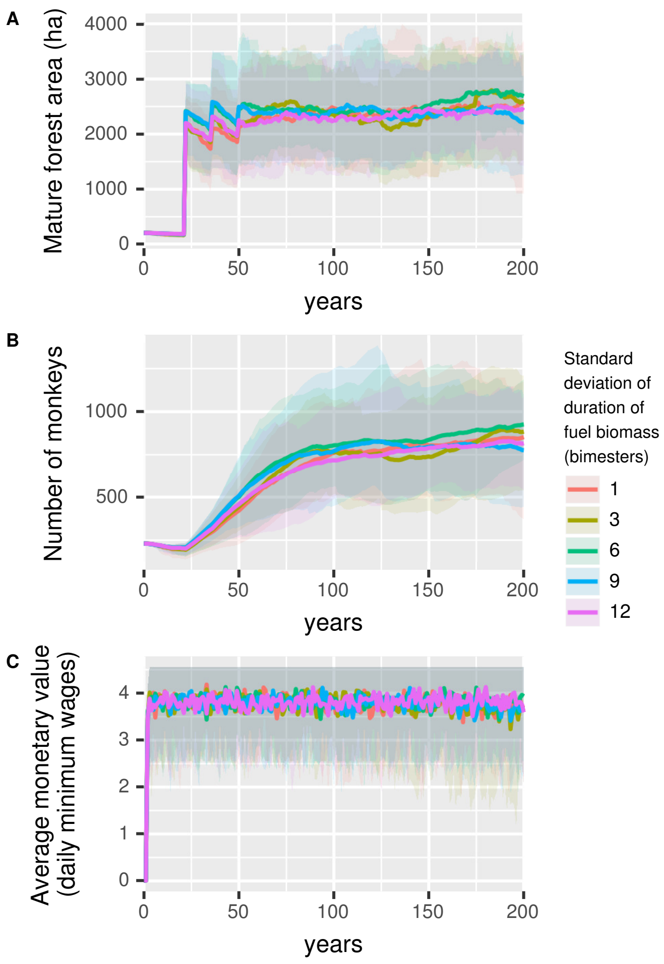


**Figure S7.** Sensitivity analysis of standard deviation of duration of fuel biomass (bimesters). Each panel represents the change in an output variable for a 200 year period and 30 simulation runs for each value of the standard deviation of duration of fuel biomass. A): Mature forest area in ha (forest with successional age greater than 50 years). B): total number of monkeys. C): average monetary value of activities done by the households in daily minimum wages. Lines represent the mean and upper and lower shadows represent 5th and 95th quartiles, respectively, for each value of the standard deviation of duration of fuel biomass (different colors).
